# Novel regimens of bedaquiline-pyrazinamide combined with moxifloxacin, rifabutin, delamanid and/or OPC-167832 in murine tuberculosis models

**DOI:** 10.1101/2021.12.21.473769

**Authors:** Rokeya Tasneen, Andrew Garcia, Paul J. Converse, Matthew D. Zimmerman, Veronique Dartois, Ekaterina Kurbatova, Andrew A. Vernon, Wendy Carr, Jason E. Stout, Kelly E. Dooley, Eric L. Nuermberger

## Abstract

A recent landmark trial showed a 4-month regimen of rifapentine, pyrazinamide, moxifloxacin and isoniazid (PZMH) to be non-inferior to the 6-month standard of care. Here, two murine models of tuberculosis were used to test whether novel regimens replacing rifapentine and isoniazid with bedaquiline and another drug would maintain or increase the sterilizing activity of the regimen. In BALB/c mice, replacing rifapentine in the PZM backbone with bedaquiline (i.e., BZM) significantly reduced both lung CFU counts after 1 month and the proportion of mice relapsing within 3 months after completing 1.5 months of treatment. Addition of rifabutin to BZM (BZMRb) further increased the sterilizing activity. In the C3HeB/FeJ mouse model characterized by caseating lung lesions, treatment with BZMRb resulted in significantly fewer relapses than PZMH after 2 months of treatment. A regimen combining the new DprE1 inhibitor OPC-167832 and delamanid (BZOD) also had superior bactericidal and sterilizing activity compared to PZM in BALB/c mice and was similar in efficacy to PZMH in C3HeB/FeJ mice. Thus, BZM represents a promising backbone for treatment-shortening regimens. Given the prohibitive drug-drug interactions between bedaquiline and rifampin or rifapentine, the BZMRb regimen represents the best opportunity to combine, in one regimen, the treatment-shortening potential of the rifamycin class with that of BZM and deserves high priority for evaluation in clinical trials. Other 4-drug BZM-based regimens and BZOD represent promising opportunities for extending the spectrum of treatment-shortening regimens to rifamycin- and fluoroquinolone-resistant tuberculosis.

## INTRODUCTION

Although effective treatment for drug-susceptible (DS) tuberculosis (TB) exists, development of new multidrug regimens capable of shortening the duration of treatment should reduce the logistical challenges of providing supervised therapy and increase treatment completion rates. Such potent regimens may have additional salutary benefits such as improving cure rates among those that do not complete treatment or have imperfect adherence. To the extent they are not compromised by resistance to isoniazid and/or rifampin, they also may extend the spectrum of activity to include isoniazid- and rifamycin-monoresistant, as well as multidrug-resistant (MDR), forms of TB, which are currently associated with worse outcomes than first-line treatment of DS-TB (1, 2).

The 3-drug combination of bedaquiline, pyrazinamide and moxifloxacin (BZM) has exhibited bactericidal and sterilizing activity superior to that of the first-line regimen of rifampin, pyrazinamide and isoniazid (RZH), with or without ethambutol (E), in multiple murine TB models, including high-dose aerosol and intravenous infection models in immunocompetent mice, aerosol infection models in immunocompromised athymic nude mice, and in C3HeB/FeJ mice that develop large caseating lung lesions (3-7). In a high-dose aerosol infection model, the sterilizing activity of BZM was also superior to that of the rifapentine, pyrazinamide and moxifloxacin (PZM) combination (6), while in a high-dose intravenous infection model, the bactericidal activity was superior to that of PZM but the sterilizing activity could not be clearly differentiated (3). As PZM forms the core of the 4-month PZMH regimen recently shown to be non-inferior to the 6-month first-line regimen in the landmark trial Tuberculosis Trials Consortium (TBTC) Study 31/ AIDS Clinical Trials Group (ACTG) A5349 (ClinicalTrials.gov Identifier: NCT02410772) (8), the results in mice suggest that BZM-based regimens may also have the potential to shorten TB treatment to 4 months or less.

The addition of the nitroimidazole prodrug pretomanid (Pa) to the BZM combination further increases the regimen’s efficacy in 3 of the murine models mentioned above (7). Moreover, in the phase 2b NC-005 trial, BZMPa produced faster sputum culture conversion in patients with MDR-TB than RZHE produced in patients with DS-TB (9) and it is now being evaluated as a 4-month regimen in the SimpliciTB trial (ClinicalTrials.gov Identifier: NCT03338621). To our knowledge, the contribution of the other marketed nitroimidazole pro-drug delamanid (D) to BZM has not been investigated. At doses judged to produce human-equivalent exposures, delamanid has exhibited comparable activity to pretomanid in BALB/c mice and in NOS2-deficient mice that develop necrotic lung lesions (10, 11) making the BZMD regimen worthy of study in pre-clinical TB models.

The rifamycin class of TB drugs has strong, exposure-dependent sterilizing activity. In murine TB models, the addition of rifampin or rifapentine further increases the strong bactericidal and sterilizing activity of combinations containing BZ (3, 6, 12, 13). For example, the combinations of BZP and BZMP are more effective than BZ and BZM, respectively (3, 6, 13). However, there is reluctance to evaluate such combinations in humans due to the strong inductive effects of these rifamycins on the metabolism of bedaquiline, a cytochrome P450 3A (CYP3A) substrate (14-16). Rifabutin (Rb) results in far less metabolic induction of bedaquiline and, therefore, may be a more suitable rifamycin for use with bedaquiline (14, 15). Rifabutin also causes less induction of metabolism of many other cytochrome P450 substrates, including certain classes of anti-retroviral drugs that may be used together with TB therapy. Rifabutin at 5 mg/kg was as effective as rifampin at 10 mg/kg and approached the efficacy of rifapentine 5 mg/kg in a murine model of latent TB infection (17). However, the contribution of rifabutin to the efficacy of combination therapy including bedaquiline, and particularly BZM, has not been investigated in pre-clinical models or clinical trials.

Decaprenylphosphoryl-β-d-ribose 2′-epimerase (DprE1) is a mycobacterial enzyme required for the biosynthesis of arabinogalactan, an essential component of the bacterial cell wall. It is targeted by 4 drugs currently in Phase 2 trials, namely the benzothiazinones BTZ-043 and PBTZ169, the azaindole TBA-7371 and the carbostyril drug OPC-167832 (O) (18-21). These drugs are bactericidal in mouse TB models, but OPC-167832 was recently shown to have superior efficacy at lower doses in a head-to-head comparison in a C3HeB/FeJ mouse infection model (22). Addition of either PBTZ169 or TBA-7371 increased the bactericidal activity of bedaquiline in BALB/c mouse infection models and PBTZ169 was further shown to increase the activity of the BZ combination (18, 20). The BO combination was more active than B alone and BMDO had greater sterilizing activity than the RHZE control in mice, although the specific contribution of OPC-167832 to this 4-drug regimen was not assessed (19). To our knowledge, the contribution of O to BZM has not been investigated in an animal model. As there may be additional benefits to replacing moxifloxacin, a broad-spectrum antimicrobial that increases the risk of *Clostridioides difficile* infection and fluoroquinolone resistance among other pathogens and commensal bacteria as off-target effects, it is also worthwhile to determine whether the narrow-spectrum drug OPC-167832 could replace moxifloxacin in the BZMD regimen, as BZOD.

Evaluation of novel drug regimens in the high-dose aerosol BALB/c mouse infection model has enabled prioritization of more efficacious regimens and provided reasonable estimates of their treatment-shortening potential relative to the first-line RHZE regimen, including regimens substituting rifapentine for rifampin and/or moxifloxacin for isoniazid or ethambutol, and the more novel PaMZ, BPaMZ and BPa+linezolid regimens (23). The present study was undertaken to minimize risk in the development of a phase 2C trial protocol to study novel regimens based on a BZ backbone. We hypothesized that the addition of rifabutin, delamanid or OPC-167832 would increase the efficacy of the BZM base regimen and that the BZOD regimen would be as effective as BZMD. The first-line 2RZHE/4RH regimen and the PZM backbone of the PZMH regimen studied in Study 31/A5349 were included as controls (8). PZM was studied rather than PZMH in order to compare directly the sterilizing activity of rifapentine and bedaquiline when either is combined with moxifloxacin and pyrazinamide.

The C3HeB/FeJ mouse model enables assessment of regimen efficacy against both intracellular and extracellular tubercle bacilli in caseating lung lesions that may be more representative of caseating lung lesions in humans (24-26). Such lesions are associated with higher bacterial burdens, alterations in drug distribution into the lesions and differing microenvironments that may affect bacterial phenotypic susceptibility and drug action and provide results that are more representative of “hard-to-treat” patients who are more likely to relapse. For example, bedaquiline distributes slowly into caseum, and pyrazinamide activity against bacteria inside caseous lesions is reduced, as a result of the near neutral pH of the caseum microenvironment (24, 27). Likewise, the differential distribution of rifapentine and rifampin into caseous lesions and the reduced efficacy of pyrazinamide in such lesions may alter the contribution of these component drugs to the overall efficacy of a regimen (as compared to their observed contribution in BALB/c mice) (28-30). Regarding the latter point, the contribution of pyrazinamide to the sterilizing activity of the RZHE regimen was previously shown to extend beyond 2 months in a C3HeB/FeJ model but not a BALB/c mouse model (28). Thus, studies in C3HeB/FeJ mice may further inform decisions on translational issues such as confirming the treatment-shortening potential of BZM-containing regimens relative to RZHE and PZMH, as well as the duration of the pyrazinamide contribution to BZM-containing regimens. The contribution of pyrazinamide to BZ-containing multidrug regimens has been assessed in BALB/c, but not C3HeB/FeJ, mice (7). Therefore, we compared the BZMRb regimen to the RZHE and PZMH regimens in this model and sought to delineate whether pyrazinamide contributes activity to the BZMRb regimen beyond the first 2 months of treatment. We also tested the novel BZOD regimen.

## RESULTS

### Assessment of bactericidal activity in BALB/c mice

Using a high-dose aerosol infection model in BALB/c mice, we evaluated the efficacy of the regimens described in Table 1. Bactericidal activity was assessed on the basis of lung CFU counts after 1 month of treatment (M1). One mouse in each of the following groups died as a result of independent gavage accidents during the first week of treatment and could not be assessed at M1: BZMD, BZMO, and BZOD. Treatment with RZHE reduced the bacterial burden by more than 2.5 log_10_ CFU at M1. All other regimens were significantly more bactericidal (p<0.0001), and all BZ-containing regimens also were significantly more bactericidal than PZM (p<0.0001) at M1. There were no statistically significant differences between BZM alone and the 4-drug BZM-containing regimens, nor between BZMD and BZOD at M1.

**TABLE 1.** Lung CFU counts assessed during treatment and proportion of BALB/c mice relapsing after treatment completion

| Regimen | Mean (±SD) lung log<sub>10</sub> CFU counts<br>(number of mice assessed at M1) | | | Proportion relapsing after treatment | | | |
| --- | --- | --- | --- | --- | --- | --- | --- |
|  | D-16 | D0 | M1 | M1 (+3) | M1.5 (+3) | M3 (+3) | M4 (+3) |
| Untreated | 4.05 ± 0.10 | 7.32 ± 0.14 | | | | | |
| 2RZHE/RH | | | 4.75 ± 0.15<br>(3) | | | 15/15 | 8/15 |
| PZM | | | 3.56 ± 0.16<br>(5) | | 13/14 | 0/15 | |
| BZM | | | 1.01 ± 0.28<br>(5) | 15/15 | 9/15 | | |
| BZMRb | | | 0.90 ± 0.25<br>(5) | 15/15 | 1/15 | | |
| BZMD | | | 1.22 ± 0.32<br>(4) | 15/15 | 8/14 | | |
| BZMO | | | 0.90 ± 0.26<br>(4) | 15/15 | 9/15 | | |
| BZOD | | | 1.30 ± 0.24<br>(4) | 15/15 | 5/15 | | |
| D-17 = 1 day after aerosol infection (n = 6 mice); D0 = day of treatment initiation, 17 days after infection (n = 6 mice). M1 = treated for one month (4 weeks); M1 (+3) = treated for one month, held for 3 additional months (12 weeks) without treatment, then sacrificed to determine the proportion with relapse, M1.5 (+3) = treated for 1.5 months and held for 3 months, etc. As of D0, 15 mice were allocated for relapse assessment at each time point indicated by proportions in the table. Due to 2 unscheduled deaths occurring after D0, only 14 mice were assessable in some arms at some time points, as detailed in the text. |  |  |  |  |  |  |  |

### Assessment of sterilizing activity in BALB/c mice

Sterilizing activity was assessed on the basis of the proportion of mice relapsing after different durations of treatment. All mice relapsed within 90 days (3 months) after one month of treatment, irrespective of treatment allocation (Table 1). One mouse in the BZMD M1.5 (+3) group was found dead of unknown causes two months into the relapse follow-up period and could not be assessed for relapse. One mouse in the PZM M1.5 (+3) group died due to a gavage accident on the final day of treatment and could not be assessed for relapse. It was culture-positive at the time of death but had a low CFU count and was therefore censored because it could not be confidently assigned a relapse or cure outcome. Among mice treated for 1.5 months, the addition of rifabutin to BZM (BZMRb) significantly (p=0.005) reduced the proportion of mice relapsing compared to BZM alone. No other BZ-containing regimen was significantly different from BZM. However, all BZ-containing regimens except BZMRb, when given for 1.5 months, resulted in significantly lower rates of relapse compared to mice treated with 2RZHE/RH for 3, and similar rates of relapse compared to mice given 2RZHE/RH for 4 months. Compared to mice treated with PZM for 1.5 months, there were significantly fewer relapses among mice treated for the same duration with BZMRb (p<0.0001) or BZOD (p=0.002). Mice treated with BZMD and BZMO for 1.5 months had fewer relapses compared to mice treated with PZM for 1.5 months, but the differences did not reach statistical significance.

### Assessment of bactericidal activity in C3HeB/FeJ mice

Using a low-dose aerosol infection model in C3HeB/FeJ mice, we evaluated the efficacy of the regimens described in Table 3. As expected in this model, there was mouse-to-mouse variability in the rate and extent of disease progression. Prior to mice being allocated to treatment groups, three mice reached a humane endpoint requiring euthanasia within 7 weeks after infection and 9 additional mice had rapid disease progression as indicated by ruffled fur and a body weight below 20g before 6 weeks post-infection. The latter 9 mice were allocated to a separate “early treatment” cohort that was initiated on treatment beginning 6 weeks (40 days) post-infection and analyzed separately from the main treatment cohort. Results for these 9 mice are presented after results for the main cohort are presented below. The remaining 275 mice were included in the main cohort, which initiated treatment one week later, at 7 weeks (47 days) post-infection (D0).

In the main treatment cohort, heterogeneity in the number and size of caseating lesions was observed upon gross inspection of the lungs at D0 (Fig. S1), as previously described (24, 25). Mice with greater lung involvement had lower body weights and higher lung CFU counts. Average mouse body weights at the start of treatment were similar across the treatment arms, except that mice in the PZMH group were significantly heavier than mice in the BZOD group (p=0.0317). Five mice allocated to treatment in the main cohort did not survive to a designated relapse endpoint. One mouse in each of the PZMH and BZMRb groups died during the first week of treatment due to apparent progression of disease and a gavage accident, respectively. One mouse in the BZOD group failed to gain weight on treatment and died after 6 weeks of treatment. One mouse in the RZHE group died after 4 months of treatment with an apparent bacterial superinfection of the scalp. Finally, one mouse in the BZMRb group died for unknown reasons 2 weeks after completing 3 months of treatment. Due to these 5 unscheduled deaths, the number of mice assessed for relapse was reduced from 19 to 18 at some time points. The results of lung CFU counts after 1 month of treatment in the main cohort are presented in Table 2. Although all regimens significantly reduced the bacterial burden after one month of treatment compared to Day 0, heterogeneity in lung CFU counts was greater in C3HeB/FeJ mice compared to BALB/c mice (Figure 1). Overall, the CFU counts in the RZHE, PZMH, BZMRb and BZOD groups were not significantly different from each other after one month of treatment. However, within each treatment group, C3HeB/FeJ mice clustered into those with higher lung CFU counts, more severe lung involvement and lower body weight and those with lower CFU counts, less severe lung involvement and higher body weight (Fig. S2). Thus, the apparently dichotomous response to treatment within each arm largely reflects the heterogeneity in the rate and extent of development of large caseating lesions. The impact of the heterogeneous pathology was particularly marked in the RZHE group, with average CFU counts in the lower CFU cluster being approximately 4 log_10_ lower than those in the higher CFU cluster (Fig. 1). In both the RZHE and PZMH groups, mice in the lower CFU clusters had CFU counts similar to those observed in BALB/c mice receiving the same treatments. On the other hand, even the lowest CFU counts among C3HeB/FeJ mice receiving BZMRb and BZOD were 2-3 log_10_ higher than those in BALB/c mice receiving the same regimens. On average, at M1, C3HeB/FeJ mice receiving BZMRb or BZOD had more extensive lung lesions upon qualitative gross lung inspection and lower body weights compared to those receiving RZHE or PZMH (Fig. S3).

**TABLE 2.**
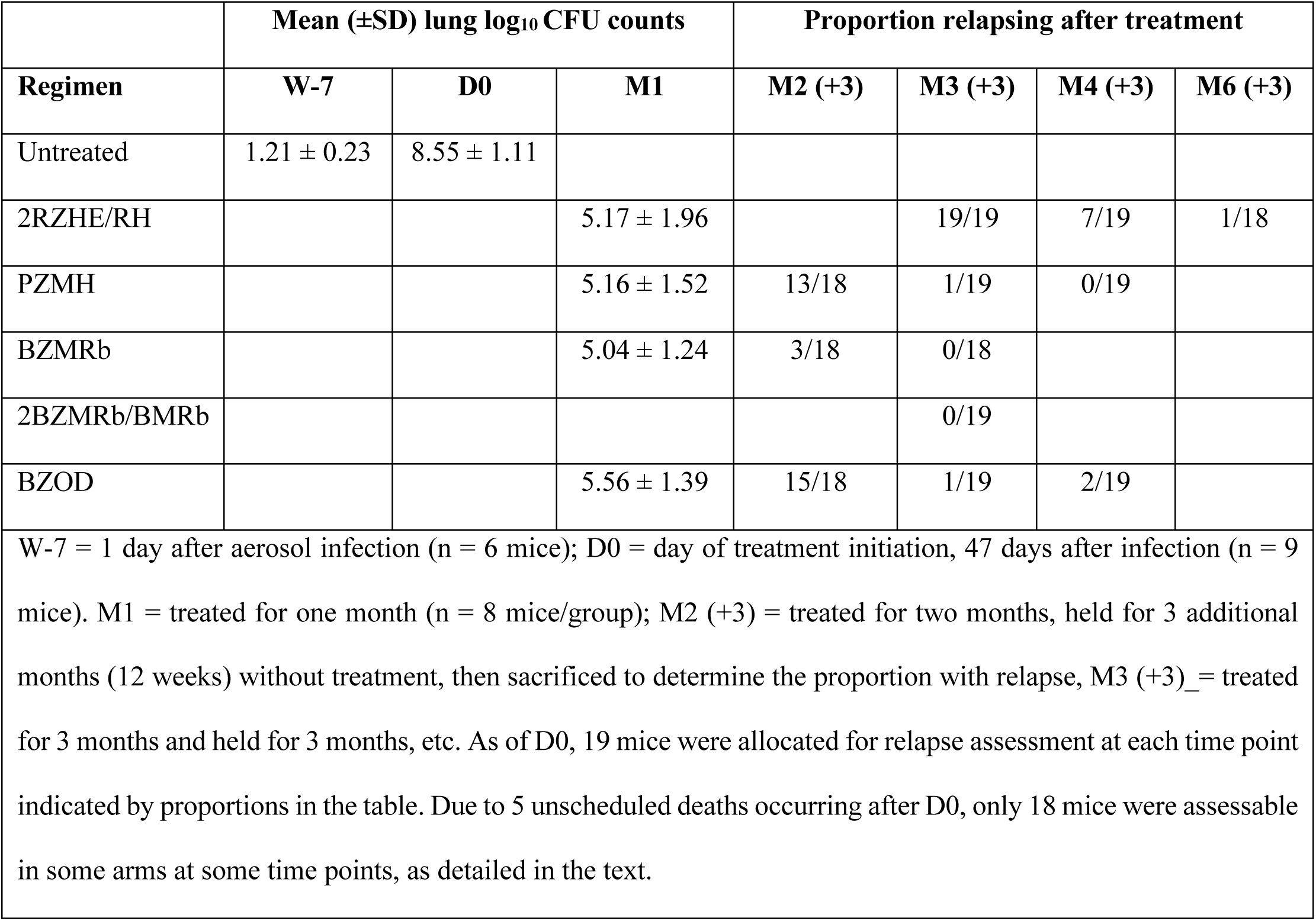
Lung CFU counts assessed during treatment and proportion of C3HeB/FeJ mice relapsing after treatment completion in the main treatment cohort

**FIG 1.**
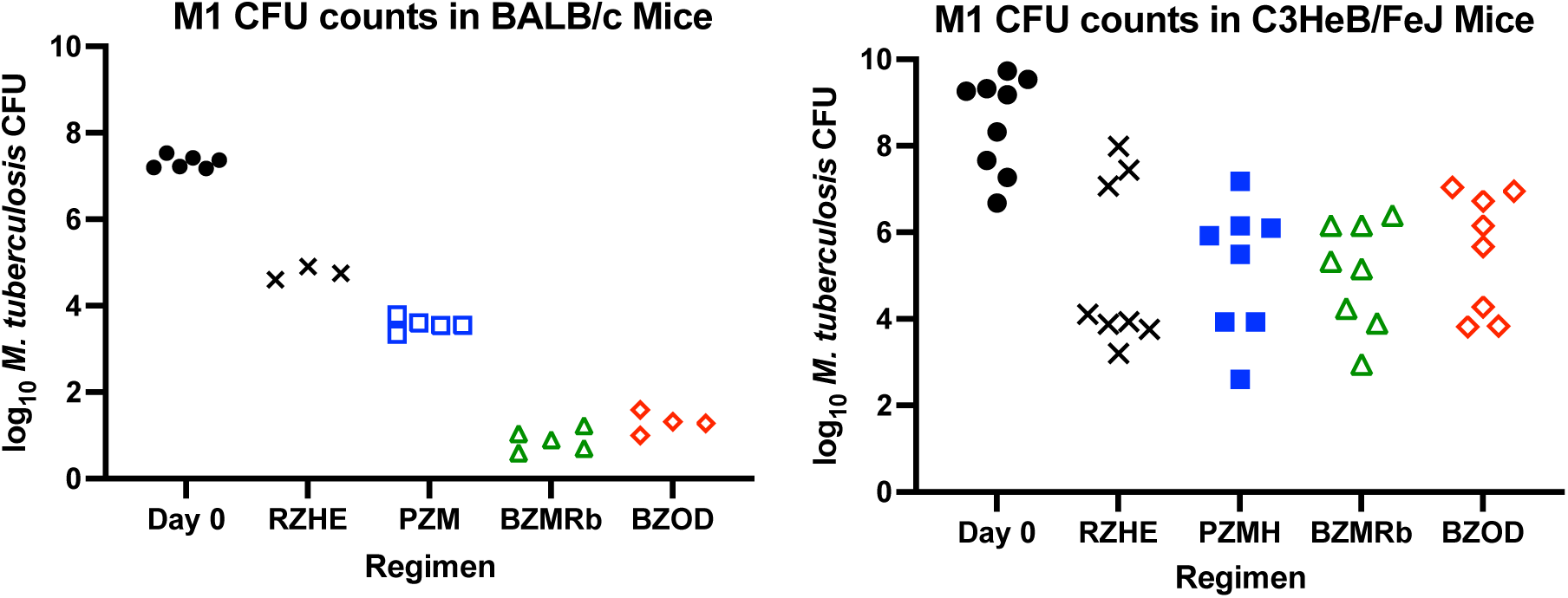
*M. tuberculosis* CFU counts in lungs of BALB/c (left) and C3HeB/FeJ (right) mice on Day 0 and after one month of treatment. All treatment regimens significantly reduced the burden compared to Day 0. In BALB/c, but not C3HeB/FeJ mice, the BZMRb and BZOD regimens were significantly more bactericidal than RZHE.

In the separately analyzed “early treatment” cohort of 9 mice, 1 mouse was sacrificed at the initiation of treatment to estimate the baseline CFU count and 4 mice each were initiated on treatment with either PZMH or BZMRb. The mouse sacrificed at treatment initiation had 9.26 log_10_ CFU in the lungs. One mouse in the PZMH group was found dead after 2 months of treatment and had 5.34 log_10_ CFU in the lungs. Two mice in the BZMRb group died after 2.5 months of treatment and had CFU counts below the limit of detection for these samples of 2.70 log_10_ CFU. The remaining 3 and 2 mice in these groups, respectively, completed 3 months of treatment and had CFU counts below the limit of detection of 5 CFU at that time point.

### Assessment of sterilizing activity in C3HeB/FeJ mice

Significant differences in sterilizing activity between some regimens emerged after completion of 2 months of treatment. Only 3 of 18 mice relapsed in the BZMRb group compared to 13 of 18 in the PZMH group and 15 of 18 in the BZOD group at this time point (p=0.002 and p=0.0002 for BZMRb vs. PZMH and BZOD, respectively) (Table 2). After 1 additional month of treatment, no mouse relapsed in the BZMRb group whether or not pyrazinamide was continued, and only 1 mouse each from the PZMH and BZOD groups relapsed. In contrast, all mice receiving RZHE relapsed. After 4 months of treatment, 7 of 19 mice from the RZHE group relapsed within 3 months, while no mouse in the PZMH group relapsed and 2 of 19 mice from the BZOD group and (p=0.008 and p=0.1245 for RZHE vs. PZMH and BZOD, respectively). Finally, after 6 months of treatment, only 1 of 19 mice in the RZHE group relapsed within 3 months after treatment.

Given the persistent small number of mice in the BZOD group relapsing after 4 months of treatment, the 2 relapse isolates from M4(+3) were tested for susceptibility to bedaquiline, pretomanid (as a surrogate for delamanid) and pyrazinamide. Both isolates were fully susceptible to these drugs.

### Pharmacokinetics of drugs in the BZOD regimen

To confirm the expected exposures of individual drugs when delamanid and OPC-167832 were co-administered with bedaquiline and pyrazinamide, plasma concentrations were measured after a single dose of BZOD in BALB/c mice. The concentrations and pharmacokinetic (PK) parameters of each drug are shown in Table S1. Plasma concentrations and PK parameters obtained following multiple doses of BZOD in infected C3HeB/FeJ mice are shown in Table S2.

## DISCUSSION

In the current study, we identified novel 4-drug BZ-containing regimens that have superior efficacy compared to RZHE and, in some cases, PZM(H). Most notably, the addition of rifabutin to BZM significantly reduced the proportion of BALB/c mice relapsing after 1.5 months of treatment and BZMRb was statistically superior to PZM. BZMRb also resulted in superior efficacy when compared to BZM using a novel pharmacodynamic biomarker, the RS ratio, in a companion study (31). Likewise, in C3HeB/FeJ mice, which develop caseating lung lesions, BZMRb had superior sterilizing activity compared to PZMH after 2 months of treatment. Although regimens combining BZ±M with rifampin or rifapentine previously were shown to be superior to RZH and even PZM in murine models (5-7), rifabutin is the only rifamycin that can be administered with bedaquiline clinically without greatly reducing bedaquiline exposures (14, 15). Therefore, the BZMRb regimen may represent the best opportunity to combine, in one regimen, the treatment-shortening potential of the rifamycin class with that of BZM and deserves high priority for evaluation in clinical trials.

In addition to having a lower risk of drug-drug interactions compared to the PZM backbone, the BZM backbone would retain efficacy against many MDR-TB strains and could therefore have an expanded spectrum of use that includes rifamycin-resistant, pyrazinamide-susceptible TB. Indeed, the ongoing SimpliciTB trial is evaluating the use of BZMPa for 4 months against DS-TB and for 6 months against TB with resistance to rifampin and/or isoniazid. In the current study, the BZMD regimen containing the other marketed nitroimidazole, delamanid, in place of pretomanid was significantly better than RZH and at least as effective as PZM in BALB/c mice, indicating that it may also be effective as a 4-month regimen. However, as a contribution of delamanid to the BZM backbone was not evident in this study, other new drugs should be evaluated in combination with BZM to identify even more effective regimens that retain activity against rifamycin-resistant TB.

Combining the DprE1 inhibitor, OPC-167832, with BZM could offer another option for both DS-TB and many MDR-TB patients. OPC-167832 is being evaluated in combination with bedaquiline and delamanid in a phase 2 early bactericidal activity (EBA) trial (NCT03678688). Like BZMD, BZMO was at least as effective as PZM in our study, although a specific contribution of O to the regimen was not observed. As the optimal clinical dose of OPC-167832 remains unsettled, it is uncertain how well the dose tested here will match the exposures that could be attained in TB patients if the drug is ultimately approved for human use. Evaluation of higher doses of OPC-167832 in mice, including C3HeB/FeJ mice (22), could be warranted, depending on the outcomes of ongoing and future clinical trials.

The novel BZOD regimen resulted in numerically fewer relapses after 1.5 months of treatment than BZM, BZMD or BZMO and significantly fewer relapses than PZM in BALB/c mice. BZOD had efficacy similar, but not superior, to PZMH in C3HeB/FeJ mice. Replacing moxifloxacin with a novel drug class in a BZM-containing regimen could have several advantages, including greater utility against fluoroquinolone-resistant strains, reduced risk of off-target effects such as selection of fluoroquinolone resistance in other pathogenic bacteria and *C. difficile* colitis. Evidence that OPC-167832 and delamanid increase the EBA of bedaquiline in the aforementioned phase 2 trial would provide further impetus to study this combination.

Our study has limitations. First, bactericidal activity was assessed following only one month of treatment in the C3HeB/FeJ mouse model. Earlier CFU assessments (e.g., at Week 2) may have revealed faster initial responses to RZHE and PZMH given the qualitatively smaller lung lesions and higher weights at M1 in these groups, while later assessments may have detected statistically significant differences in lung CFU counts that correlated with relapse outcomes. Second, quantification of drugs in plasma was not performed for all regimens. Therefore we cannot fully exclude the possibility that an unmeasured drug-drug interaction affected one or more group-wise comparisons. Finally, drug concentrations were not measured in lung lesions to determine the extent to which relative differences in regimen performance between the mouse models might be explained by differences in drug partitioning into caseating lesions.

In conclusion, novel regimens of BZM combined with rifabutin, delamanid, or OPC-167832 or the BZOD regimen display similar or better sterilizing activity compared with PZMH, a regimen recently shown to be non-inferior as a 4-month regimen when compared to the 6-month RZHE/RH regimen in a Phase 3 trial. These regimens merit further assessment in clinical trials.

## MATERIALS AND METHODS

### Murine infection models

All experimental designs and procedures were approved by the Animal Care and Use Committee of Johns Hopkins University. Female BALB/c mice (Charles River Labs) (n = 255), 5-6 wks old, were aerosol-infected in two runs using an inhalation exposure system (Glas-Col) with approximately 4 log_10_ CFU of *Mycobacterium tuberculosis* H37Rv, using a mouse-passaged, frozen aliquot that was thawed and then actively grown in culture to an optical density at 600nm (OD) of 0.8-1.0. Female C3HeB/FeJ mice (Jackson Labs) (n = 287), 10 wks old, were aerosol-infected in three runs with approximately 50 log_10_ CFU of *M. tuberculosis* HN878, using a mouse-passaged, frozen aliquot that was thawed and diluted prior to infection. Mice were block randomized by run to distribute the mice equivalently into the different treatment arms. Treatment started 17 days later (D0) in BALB/c mice. In C3HeB/FeJ mice, disease progressed more rapidly in 12 mice, of which 3 died prior to treatment allocation and 9 were allocated to an “early treatment” cohort that initiated treatment 40 days after infection. The remaining 275 C3HeB/FeJ mice comprised the main treatment cohort and initiated treatment 47 days post-infection. The number of bacteria (CFU/ml) in the culture used for infection was determined by plating serial 10-fold dilutions on 7H11 agar. Six untreated mice were sacrificed for lung CFU counts on the day after infection to determine the number of CFU implanted and 6 BALB/c mice and 9 C3HeB/FeJ mice in the main treatment cohort were sacrificed at D0 to determine the baseline bacterial burden at the start of treatment. Lungs were removed aseptically and homogenized in glass grinders. Serial 10-fold dilutions of lung homogenates were plated on 7H11 agar supplemented with 10% oleic acid-albumin-dextrose-catalase (OADC). Plates were incubated for 4 weeks before final CFU counts were determined.

### Drug preparation and administration to mice

The drug doses (in mg/kg indicated in subscripts) were R_10_, H_10_, Z_150_, E_100_, B_25_, M_100_, P_10_, Rb_10_, D_2.5_ and O_2_. R, H, Z, E, M, and B were obtained as previously described (5-7). Rb powder was purchased from Carbosynth. P tablets were purchased from the Johns Hopkins Hospital pharmacy. D and O were provided by Otsuka. R, H, Z, E, M, P and Rb were prepared in deionized sterile water, B in an acidified 20% hydroxypropyl-β-cyclodextrin (HPCD) solution, D and O in 5% gum arabic suspension after grinding in an agate mortar. P was prepared as previously described (32). All drugs were administered once daily and given 5 days per week by gavage in a volume of 0.2 ml. R, P and Rb were administered alone at least 1 hour before other drugs. B was administered alone at least 2 hours before other non-rifamycin drugs. ZH, ZHE, ZM, ZMH, ZMD, ZMO and ZOD were administered together in a single gavage. Except where indicated in Tables 1 or 3, Z was given for the entire treatment duration.

### Evaluation of efficacy in mice

Efficacy determinations in BALB/c mice were based on lung CFU counts after 1 month of treatment (n = 3 mice in the 2RZHE/RH group, 5 mice in other groups) and relapse assessments carried out 3 months after completing 1 and 1.5 months of treatment with each BZ-containing regimen, 1.5 and 3 months of PZM, or 3 and 4 months of 2RZHE/RH (n = 15 mice/group/time point). Efficacy determinations in C3HeB/FeJ mice were based on lung CFU counts after 1 month of treatment (n = 8 mice/group) and relapse assessments carried out 3 months after completing 2 and 3 months of treatment with BZMRb, 2, 3 and 4 months of treatment with BZOD and PZMH, or 3, 4 and 6 months of 2RZHE/RH (n = 20 mice/group/time point intended). The time points for relapse assessments in both models were selected based on prior experience with the same or similar regimens in order to optimize the discrimination between regimens. Lung homogenates were plated in serial 10-fold dilutions on 7H11 plates supplemented with 10% OADC and 0.4% activated charcoal to reduce drug carryover. For M1 and later time points, plates were incubated for 6 weeks before final CFU counts were determined. Relapse was assessed by plating the entire lung homogenate. Mice were considered to have relapsed if growth of a single colony or more was detected.

### Evaluation of selected relapse isolates for drug resistance

An isolate from each of the 2 C3HeB/FeJ mice that relapsed after 4 months of treatment with BZOD were tested for susceptibility to bedaquiline, pretomanid and pyrazinamide. The isolates were collected by scraping together colonies from one agar plate used to test for relapse at M4+3, suspending them in phosphate-buffered saline (PBS) and homogenizing with glass beads using a bead beater. The resulting suspension was left to settle for 30 min before the supernatant was removed and plated in serial 10-fold dilutions on drug-free 7H11 plates and 7H11 plates containing either 0.125 µg/ml of bedaquiline or 2 µg/ml of pretomanid, as previously described (7, 11). The HN878 stock strain was grown in 7H9 broth to an OD of approximately 1 and plated in the same fashion to serve as a control. Resistance was defined as a CFU count on drug-containing plates that was ≥1% of the CFU count observed on drug-free plates.

An aliquot of each suspension containing approximately 10^6^ CFU was used to determine the pyrazinamide MIC using the broth macrodilution method. Briefly, polystyrene tubes containing 7H9 broth supplemented with 10% OADC were prepared with serial 2-fold dilutions of pyrazinamide ranging from 75 to 600 µg/ml, as well as a tube containing 900 µg/ml pyrazinamide and a drug-free control tube. Each tube was inoculated with approximately 10^5^ CFU/ml of either relapse isolate or the HN878 stock strain as control. MIC was defined as the lowest concentration preventing visible growth after 14 days of incubation at 37°C. Resistance was defined as an MIC higher than the MIC against the HN878 stock strain. Pyrazinamidase activity was also determined for each isolate using a previously described method (33).

### Plasma PK analysis

To confirm the expected exposures of B, Z, O and D when administered as BZOD, uninfected BALB/c mice received a single dose of each drug in combination according to the dosing schedule of the main efficacy experiment (B administered 2 h prior to ZOD). Approximately 50 µl of whole blood was collected in-life by mandibular puncture at 3, 4, 6 and 9 h and by cardiac puncture under isoflurane anesthesia at 25 (24+1) h after dosing of the B component. Three mice were sampled per time point. Each mouse was sampled only once or twice prior to the terminal time point. Blood was collected into EDTA-containing tubes and processed to obtain approximately 25 µl of plasma. Blood was obtained from infected C3HeB/FeJ mice in the main efficacy experiment after 3 months of treatment. Three mice per group per time point were sampled by submandibular bleed at 3, 4, 6, 9 and 25 hours after treatment. The plasma samples were frozen at -80°C and sent to the Center for Discovery and Innovation (Hackensack Meridian Health, Nutley, NJ 07110) for quantitative analysis and pharmacokinetic modeling. Concentrations of B and its primary N-desmethyl metabolite (M2), Z, O and D in mouse plasma were determined by validated LC-MS assays (22, 34, 35). AUC_0-24h_ was determined by standard non-compartmental techniques. AUC_0-9h_ was determined for B and the M2 metabolite in BALB/c mice because a second B dose preceded sampling at the 25 hr time point.

### Statistical analyses

Group mean CFU counts were compared to those of RZHE, PZM or BZM controls using one-way ANOVA with Dunnett’s post-test (GraphPad Prism 9). Group means of the BZMD and BZOD groups were compared by t-test. Differences in relapse proportions were assessed by Fisher’s Exact test using the Holm-Bonferroni correction for multiple comparisons. Corrected p values < 0.05 were used to define statistical significance.

## ACKNOWLEDGMENTS

This study was funded by the U.S. Centers for Disease Control and Prevention’s Antibiotic Resistance Solutions Initiative through a subcontract from Westat No. 8758-S01. Otsuka Pharmaceutical donated delamanid and OPC-167832.

## Disclaimer

The findings and conclusions in this report are those of the authors and do not necessarily represent the official position of the Centers for Disease Control and Prevention (CDC) or the U.S. Department of Health and Human Services.

